# The effect of scanning speed on texture-elicited vibrations

**DOI:** 10.1101/680322

**Authors:** Charles M. Greenspon, Kristine R. McLellan, Justin D. Lieber, Sliman J. Bensmaia

## Abstract

To sense the texture of a surface, we run our fingers across it, which leads to the elicitation of skin vibrations that depend both on the surface and on exploratory parameters, particularly scanning speed. The transduction and processing of these vibrations mediates the ability to discern fine surface features. In the present study, we seek to characterize the effect of changes in scanning speed on texture-elicited vibrations to better understand how the exploratory strategy shapes the neuronal representation of texture. To this end, we scanned a variety of textures across the fingertip of human participants at a variety of speeds (10 – 160 mm/s) while measuring the resulting vibrations using a laser Doppler vibrometer. We found that increases in speed led to systematic increases in vibratory intensity and to a systematic upward multiplicative shift in the frequency composition of the vibrations. Furthermore, we showed that the upward shift in frequency composition accounts for the increase in intensity. The enhancement of higher frequency components accounts for the observed increase in the firing rates of nerve fibers, particularly Pacinian corpuscle-associated fibers, which are most sensitive at the high frequencies.

## INTRODUCTION

To discern the texture of a surface, we spontaneously run our fingers across it (1), an exploratory procedure that results in the elicitation of skin vibrations that reflect the microstructure of the surface. While coarse textural features can be sensed without movement, our perception of fine textural features relies on the processing of skin vibrations elicited during scanning (2–5). Two populations of tactile nerve fibers – rapidly adapting (RA) and Pacinian (PC) fibers – are exquisitely sensitive to skin vibrations and produce millisecond-precision temporal spiking sequences that reflect the vibrations (3,4,6,7). These vibration-sensitive afferents mediate our ability to perceive fine textural features as evidenced by the fact that desensitizing them impairs the perception of fine texture (5).

Texture-elicited vibrations are not only sensitive to surface microstructure but also to exploratory parameters, particularly scanning speed. Indeed, skin vibrations dilate or contract with decreases or increases in scanning speed (7,8), respectively, resulting in concomitant dilations or contractions of the evoked spiking sequences in the nerve (6). In addition, the firing rates of nerve fibers and of their downstream targets tend to increase with increases in scanning speed (9,10). Whether this enhanced neural response reflects the contraction of the spike trains or is caused by an increase in the intensity of the vibrations – which would also lead to higher firing rates – remains to be elucidated.

Previous measurements of the effect of scanning speed on texture-elicited vibrations were obtained with textures scanned at a single speed through each experimental block (7) and, given the measurement approach, the overall gain of the measurements varied from block to block. As a result, the raw magnitude of the vibrations could not be compared across experimental blocks and so the effect of speed on vibratory intensity could not be assessed. To fill this gap, we used a laser Doppler vibrometer to measure the vibrations evoked in the skin when everyday textures are scanned across the skin over a range of behaviorally relevant speeds within single blocks (11).

We found (as expected) that the frequency composition of vibrations shifts to higher or lower frequencies with increases or decreases in scanning speed, respectively. Furthermore, vibratory intensity increases with speed but does so in a texture-specific way. We then demonstrated that changes in vibratory intensity with scanning speed can be explained by the multiplicative shift in the frequency composition of the vibrations. We discuss the implications of these findings for the neural coding of both texture and speed.

## RESULTS

With a custom-built rotating drum stimulator (see ref. (7)), we scanned 8 textured surfaces across the right index fingertip of 5 human subjects (2 male, 3 female, age range 20-25 years) at 28 speeds spanning the range used in natural texture exploration (ranging from 10 to 160 mm/s)(11) while measuring the evoked skin vibrations using a laser Doppler vibrometer (Polytec OFV-3001 with OFV 311 sensor head; Polytec, Irvine, CA).

### Effect of scanning speed on the intensity of texture-elicited vibrations

First, we found that the intensity of the vibrations elicited in the skin, as indexed by RMS vibratory speed (VS_RMS_), varied widely across textures (**Figure 1**)(3-way ANOVA, F(7,1117) = 1477.83, p < 0.001, η^2^ = 0.5189), as has been previously shown (7). Indeed, VS_RMS_ varied over more than an order of magnitude at both the slowest scanning speed (0.511 to 13.3 at 10 mm/s) and the fastest one (2.08 and 33.2 mm/s at 160 mm/s). Second, we found that VS_RMS_ increased with the scanning speed (F(27,1117) = 125, p < 0.001, η^2^ = 0.1693). The effect of speed on VS_RMS_ was texture-dependent, as evidenced by a significant texture x speed interaction (F(189,1117) = 4.37, p < 0.001, η^2^ = 0.0414). Indeed, VS_RMS_ increased two- to eleven-fold from the lowest to the highest scanning speed, depending on the texture. VS_RMS_ did not just vary across textures but also across participants (F(4,1117) = 563.53, p < 0.001, η^2^ = 0.1131) and the participant x speed interaction was weak but statistically significant (F(108,1117) = 2.5, p < 0.001, η^2^ = 0.0135). We found that the relationship between VS_RMS_ and scanning speed was well described by a power law, with a different set of parameters – exponent and scaling factor – for each texture and participant (**Figure 2**).

**Figure 1.**
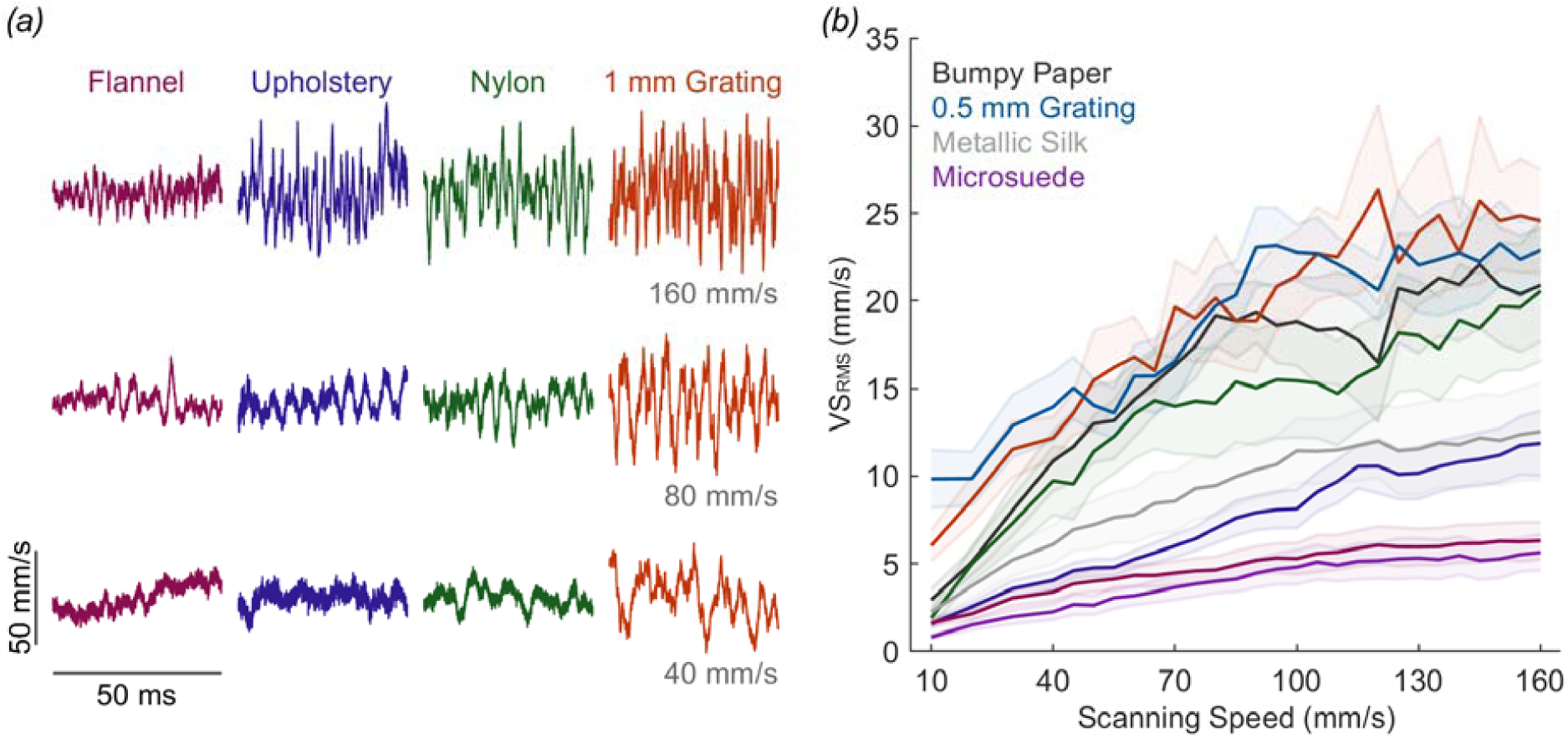
The RMS speed of texture-elicited vibrations increases as scanning speed increases. (*a*) Example velocity traces for 4 textures scanned at 3 speeds. (*b*) VS_RMS_ elicited by each texture vs. speed – solid line and shaded area indicate mean and standard error respectively.

**Figure 2.**
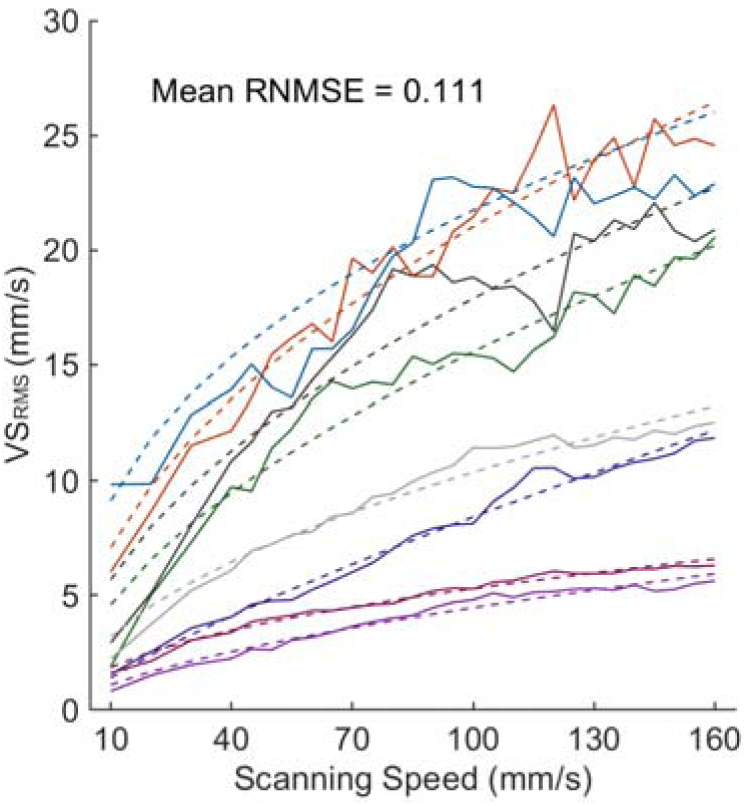
Mean measured VS_RMS_ (solid line) and the power model prediction (dashed line). Each trace denotes a different texture, following the color-scheme from **Figure 1**. For all textures and participants, fitted exponents were less than 1.

Whereas VS_RMS_ systematically increased with speed, the RMS skin displacement did not (**Supplementary Figure 1**).

### Effect of scanning speed on the frequency composition of texture-elicited vibrations

Next, we examined how the frequency composition of the vibrations changed with scanning speed. We observed that, consistent with previous findings, the power spectral density (PSD) shifted systematically towards higher frequencies as scanning speed increased (**Figure 3a**). While the spectral structure of these vibrations was relatively consistent across speeds when expressed in spatial units (by dividing the temporal frequency by the speed - **Figure 3b**)(cf. refs (7,8,12)), it was not completely speed independent. In some cases, the power at the peak frequency of a periodic texture increased with speed (**Figure 3a**, metallic silk) and, in others, it exhibited a non-monotonic relationship with speed (**Figure 3a**., microsuede).

**Figure 3.**
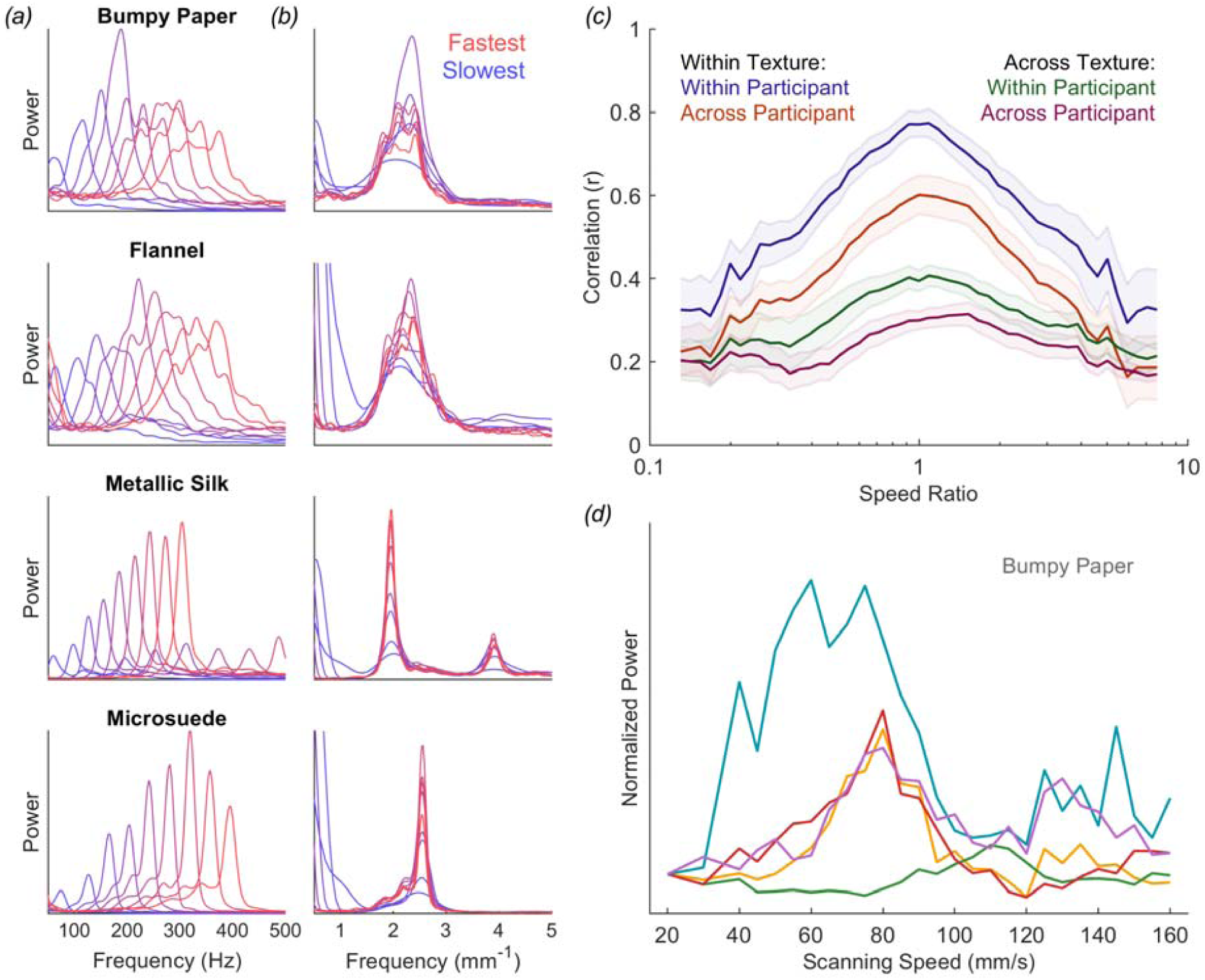
Spectral characteristics of the vibrations are shifted in the temporal domain but conserved in the spatial domain. (*a,b*) The frequency composition of the vibrations shifts to higher frequencies with increases in scanning speed but remains relatively consistent when expressed in spatial units. (*c*) Correlations within or across participants and/or textures with respect to the ratio between scanning speeds. Correlations drop as the difference in speed increases. (*d*) The relationship between peak power (normalized to that at 20 mm/s) and scanning speed for Bumpy Paper varies widely across

To quantify how the frequency composition of each texture – expressed in spatial units – varied across speed, we computed the correlation of the PSDs across all speeds within textures and participants. First, we found that the correlation across repeated presentations of the same texture was very high (mean ± s.e.m: r=0.77±0.08), emphasizing the repeatability of the measurements themselves (**Figure 3c, Supplementary Figure 2a**). Second, PSDs became more dissimilar as the difference in speed increased. At large differences in speed, the (spatial) PSDs at the two speeds for a given texture were nearly as dissimilar as were corresponding PSDs for different textures. Third, we found that the PSDs for a given texture were similar across subjects and exhibited the same drop off as speed difference increased (also see **Supplementary Figure 2b**). Comparisons of PSDs across textures revealed subject-specific structure in the vibrations as well as coincidental similarities in the vibrations elicited by certain pairs of surfaces.

Next, we examined the speed-related distortions in the PSDs. To this end, we examined the relationship between speed and the power at the peak frequency. We found that the effect of speed on the magnitude of that peak varied widely across textures for a given participant, and across participants for a given texture (see **Figure 3d** for an example, **Supplementary Figure 2c**). Speed-related distortions in the PSDs of the texture-elicited vibrations are thus highly idiosyncratic across both textures and participants.

### Speed-dependent changes in frequency drive changes in vibratory intensity

While (spatial) PSDs change with scanning speed, the effect of scanning speed on VS_RMS_ is highly systematic for all textures and subjects. With this in mind, we investigated the degree to which the speed-dependent increase in vibratory amplitude could be explained by the observed speed-dependent warping of the frequency composition of the vibrations. Accordingly, we warped the PSDs evoked by each texture at an intermediate scanning speed (90 mm/s) to that expected at the target speed. So, for example, the PSD at 45 mm/s would be the same as the vibration at 90 mm/s but shifted toward lower frequencies (**Figure 4a**). If the main effect of speed is to dilate or contract the skin vibrations, then the resulting warped PSDs would match the measured ones and the predicted VS_RMS_ would match its measured counterpart. We found that this simple model captured the relationship between VS_RMS_ and speed with remarkable accuracy (mean RNMSE= 0.131, **Figure 4b** and **Supplementary Figure 3**), despite the aforementioned idiosyncratic speed-dependent distortions in the PSDs themselves.

**Figure 4.**
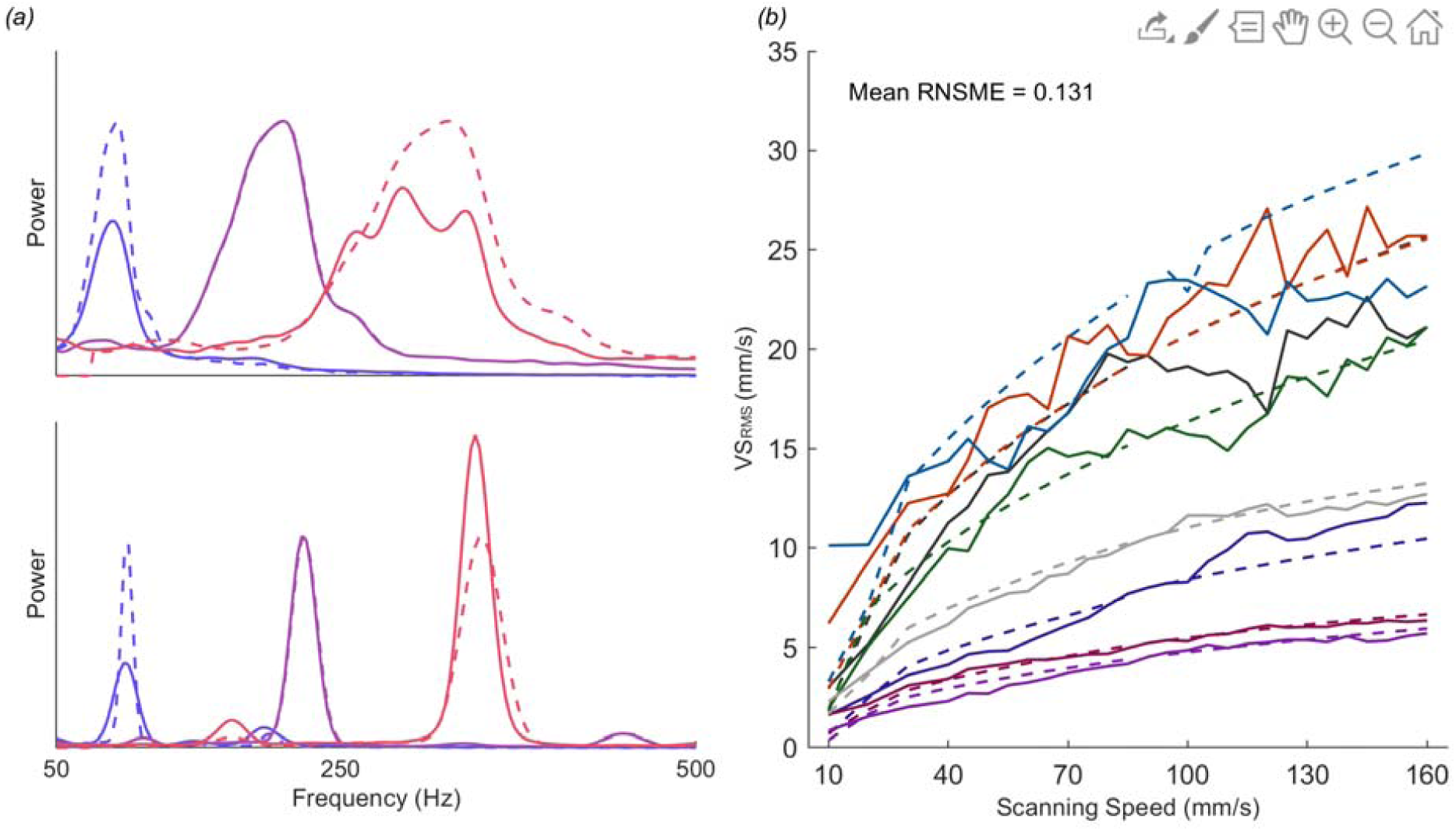
Dilations or contractions of the vibrations accounts for the effect of speed on vibratory amplitude. (*a*) Solid lines show PSDs at 3 speeds (40, 90, and 140 mm/s) for two textures, dashed lines show the PSDs obtained by warping the PSD of the 90 mm/s trace. At higher speeds, traces become wider and contribute more spectral power. (*b*) Recorded and predicted velocity based on warping the 90 mm/s trace, predictions again displayed as dashed lines (reference speed is excluded). See 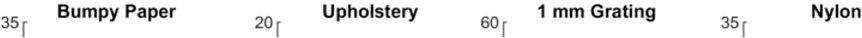

In fact, the parameter-free model performed nearly as well as did the power model, which entails 2 parameters for each texture pair and participant (**Figure 4b**). Predictions of the warping model were robust to changes in the choice of reference speed though there was a tendency for prediction error to increase at lower speeds. To ensure that this result was not an artifact of the texture set, we replicated the warping analysis with vibrometry measurements from 11 other textures scanned across the skin at 5 speeds (20, 40, 80, 120, and 160 mm/s) and obtained the same result (**Supplementary Figure 4**).

Note that this model assumes that the sole effect of scanning speed is to multiplicatively shift the PSD. As mentioned above, however, speed exerts an effect on the shape of the PSD that is idiosyncratic across textures and participants, which likely reflects idiosyncratic interactions between skin and surface that cannot be captured without a much more complex model of the biomechanical interaction. Our results suggest that these effects only have a minor impact on VS_RMS_ and thus do not lead to systematic discrepancies in the model predictions.

## DISCUSSION

### Variability across participants

While texture-elicited vibrations were similar across participants, they were not identical. Indeed, the speed-dependence of the shape of the PSDs varied widely across participants and differences across participants accounted for over 10% of the variance in the VS_RMS_. These differences are likely due to two factors. The first is that skin lubrication, which has been shown to strongly modulate the frictional interactions between skin and texture, varies widely across individuals (13,14). We attempted to minimize variations in lubrication by cleaning fingertips with an alcohol wipe, an intervention that is unlikely to have completely eliminated differences in lubrication. The second contributor to differences across subjects is variation in the biomechanics and microstructure of the skin. The mechanical properties of the skin – stiffness and elasticity – have been shown to vary substantially across individuals and impact skin-surface interactions (15–17). Similarly, fingerprint microgeometry famously differs across individuals and impacts the skin response to scanned textures (7,18–20). These various factors together combine to give rise to subject-specific interactions between and surfaces, which are then reflected in the skin vibrations. Nonetheless, the simple, parameter-free warping model accounts for the effect of scanning on VS_RMS_ with remarkable accuracy.

### Implications for texture perception

Texture perception is highly independent of scanning speed and contact force despite the fact that the response of the skin and that of tactile nerve fibers is highly dependent on these exploratory parameters (6,21). Indeed, the firing rates of tactile nerve fibers, particularly PC fibers, increase with increases in scanning speed. Here, we show that the effect of speed on afferent responses is mediated by an increase in VS_RMS_ (cf. ref. (3)), itself caused by a shift in the frequency composition of the vibrations to higher frequencies, and not changes in displacement. Both the increase in VS_RMS_ of texture-evoked vibrations and the shift to higher frequencies explains why the effect of speed is most pronounced in the responses of PC fibers, weaker in RA fibers, and weakest in SA1 fibers (22). Indeed, PC fibers are sensitive to the rate of displacement of the skin (23) and peak in sensitivity at the high frequencies (> 100 Hz)(24); RA fibers are also sensitive to vibratory speed but peak in sensitivity at intermediate frequencies; SA1 fibers are least sensitive to vibratory speed and peak in sensitivity at low frequencies. While the peripheral representation of texture is highly speed-dependent, afferent signals are differentiated downstream – spatially and temporally – and these neural computations give rise to a more speed-independent representation of texture (25).

### Implications to speed perception

While we have a sense of how fast a surface is moving across our skin (22,26), tactile speed perception is powerfully biased by surface texture: coarser textures are systematically perceived as moving faster. That speed perception is not veridical can be explained by its neural basis. Indeed, perceived speed is determined by the strength of the response evoked in PC fibers, itself dependent on both texture and scanning speed. In the present study, we replicate the finding that vibratory intensity – gauged with VS_RMS_ – is strongly dependent on surface texture. We also show that VS_RMS_ is strongly dependent on speed. Given that VS_RMS_ at the high frequencies (> 50 Hz) is a good proxy for PC firing rates, the present results account for the dependence of perceived speed on both speed and texture (22).

### Artificial textures

The rise of mobile devices – tablets and phones – has spurred the development of haptic interfaces to provide tactile feedback during manual interactions with the devices, most commonly in the form of vibrations triggered by contact with the screen. That texture perception relies in part on vibration has engendered attempts to produce elicit percepts by generating texture-like vibrations on the surface (27–32). Because texture-elicited vibrations depend on both the texture and the exploratory parameters, a challenge in generating artificial textures has been to store the information necessary to replay vibrations appropriate for a given texture and set of exploratory parameters. One approach consists in tiling the space of possible scanning speeds and contact forces and interpolating between these measured values (31). While humans naturally vary their contact force with speed (33), responses of primary afferents to textures are relatively robust to changes in force (21). Thus, a more parsimonious approach would be to store the vibratory profile of each texture at some intermediate speed (say 90 mm/s) and warping said trace depending on the instantaneous scanning speed, ignoring the relatively subtle effects of contact force on vibrations. The resulting vibrations would largely match their natural counterparts while relying on a single stored trace. While this strategy will accurately account for the effect of speed on vibratory power, however, it ignores the idiosyncratic texture- and subject-specific effects of speed on the spectral signature of the vibrations. The biomechanical determinants of these idiosyncrasies remain to be determined as does the degree to which they shape the evoked percept.

## METHODS

### Participants

Five subjects (2 male, 3 female, 20-25 years of age) participated in the main study. Another ten subjects participated in the replication study (6 male, 4 female, 19-29 years of age), the results of which are shown in Supplementary Figure 4. Procedures were approved by the Institutional Review Board of the University of Chicago.

### Data Collection

#### Texture apparatus and stimuli

The subject sat with arm resting on a padded frame. The hand was strapped to a custom hand holder with the palm facing upward and the index finger propped at a 45-degree angle. The fingertip was cleaned with alcohol to minimize variations in skin moisture across participants. On each trial, a rotating drum stimulator (previously described in (7)) scanned one of 8 or 11 texture strips (2.5 cm x 16 cm) across the subject’s fingertip at a pre-calibrated contact force (30 g) and at one of 28 speeds, ranging from 10 to 160 mm/s, evenly spaced. Trial duration ranged from 0.4 seconds at the fastest speed to 1.6 seconds at the slowest one and the inter-trial duration was 3.5 s.

#### Vibrometer

A laser-Doppler vibrometer (Polytec OFV-3001 with OFV 311 sensor head; Polytec, Irvine, CA) was used to record the vibrations elicited in the skin (cf. refs. (7,34)). Briefly, the vibrometer measures the time-varying velocity along the axis perpendicular to the surface of the skin, without making contact with it. A small strip of white-out correction fluid (BIC USA, Shelton, CT, USA) was applied to the distal aspect of the distal interphalangeal joint of the right index finger – near the location where the finger made contact with the surface – to improve the reflectance of the finger and reduce dropouts. The vibrometer beam was then directed and focused on the white-out.

### Data Analysis

#### Preprocessing

Only the last 350 ms (35,000 data points) of each trace was used to exclude transients caused by the establishment of contact with the surface and ensure that the frequency resolution of the power spectra was equivalent across speeds. For each trace, outliers – which fell 6 SD from the mean for that speed/texture pair – were replaced by linear interpolation of the two adjacent time points. Dropout-removal was performed iteratively until all were removed. Traces that contained 2 ms or more of dropouts were eliminated. Vibrations decay as they travel from their origin and do so in a frequency dependent way (see ref. (34)). We corrected for this decay based on previous measurements and assuming a distance of 1 cm between measurement location and the nearest contact between skin and surface.

#### Spectral analysis

Power spectral densities were computed using Welch’s method over the range from 50 and 1000 Hz (see below). To convert frequency spectra to the spatial domain, we divided the frequency by the scanning speed (mm/s). To compute displacement traces, each component of the Fourier spectrum was divided by 2πfj and the resulting spectra were converted back to the time domain. The RMS of the trace within the desired frequency band was then determined by:

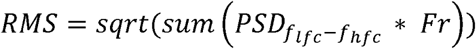

Where PSD is the power spectral density, *f* is the frequency, *lfc/hfc* are the frequency cutoffs, and *Fr* the frequency resolution of the PSD.

#### Noise filtering

We wished to determine the range over which measured vibrations are texture specific. To this end, we first normalized each PSD so that it summed to 1 and calculated the mean normalized PSD across textures, speeds, and participants (**Supplementary Figure 5a**). We observed that the bulk of the vibratory energy was contained in the first 50 Hz. Next, we examined the degree to which the power in different frequency bands varied across textures (**Supplementary Figure 5b**) and found that the low frequencies (up to ∼50 Hz) were relatively texture-independent, consistent with previous observations (7). Finally, we performed the same analysis while varying both the low- and high-frequency cutoffs (**Supplementary Figure 5c**) and found that performance was stable with high frequency cutoffs above 300 Hz. Consequently, we used 50 Hz as the low frequency cutoff and 1000 Hz as the high frequency cutoff to ensure that no texture-dependent signals were discarded.

#### Spatial PSD correlation

To directly compare the PSDs at different speeds, we expressed them in spatial units by dividing frequency by speed then resampling the PSDs via linear interpolation to achieve the same frequency resolution (matched to that of the reference speed). We then computed the correlations of pairs of PSDs (within texture/within participant, within texture/across participants, etc.)

#### Regression

We implemented three models to relate *VS*_*RMS*_ to scanning speed: linear (*VS*_*RMS*_ =*B*\* *speed*), log-linear (*VS*_*RMS*_ = *B* * *log*(*speed*)), and power (*VS*_*RMS*_ = *B* * *speed*^*k*^) where *B* and *k* are free parameters. We did not include an intercept term as lack of movement across the skin should result in no vibration. We only report the results from the power relationship as it outperformed the other two.

#### Warping model

To predict the spectrum at speed S_warp_ from the PSD measured at speed S_ref_, we first divided each frequency of the PSD at S_ref_ by the speed – to yield a spatial PSD. Next, we resampled the resulting spectrum to match the resolution of the spatial PSD at S_warp_ then multiplied each frequency by the speed of S_warp_. To then assess the predictions of the warping model, we used the range-normalized mean squared error, given by:

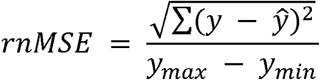

where *y* is the measured VS_RMS_ and *ŷ* is the predicted VS_RMS_. The values at the reference speed were excluded as these yielded perfect performance.

## Funding

This work was supported by National Institute of Health/National Institute of Neurological Disorders and Stroke grant RO1 NS 101325 and by NSF grant IIS-1518614 awarded to S.J.B.

## Author contributions

S.J.B., C.M.G., and J.D.L. conceived of and designed the experiment. C.M.G. and K.R.M. collected the data. C.M.G. analyzed the data. C.M.G. and S.J.B. wrote the manuscript while J.D.L. provided critical feedback.

## SUPPLEMENTARY FIGURES

**Supplementary Figure 1.**
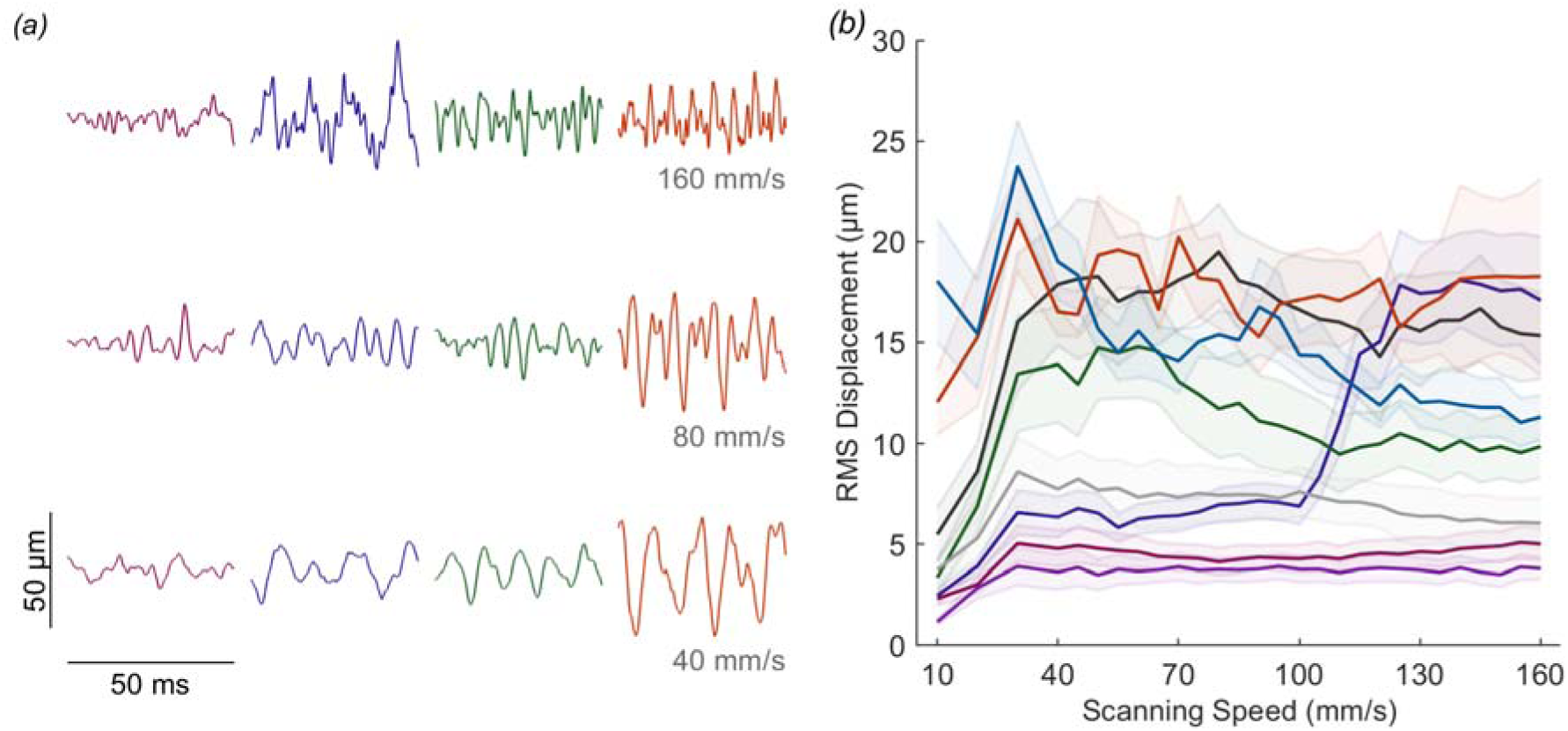
Skin displacement over time. (*a*) Example displacement traces for 4 textures scanned across the skin at 3 speeds. (*b*) RMS amplitude averaged across subjects vs. speed. A 3-way ANOVA revealed that texture is the dominant source of variance (F(7,1117) = 794.64, p < 0.001, η2 = 0.5450), followed by the participant (F(4,1117) = 208.12, p < 0.001, η2 = 0.0816), with speed as a distant third (F(27,1117) = 10.53, p < 0.001, η2 = 0.0280). The participant x texture and texture x speed interactions were both significant (F(27,1118) = 50.97, p < 0.001, η2 = 0.1398; F(189,1117) = 6.52, p < 0.001, η2 = 0.1208) while the participant x speed interaction was not (F(108,1117) = 0.87, p = 0.8223, η2 = 0.0092). Color-scheme follows that of **Figure 1**.

**Supplementary Figure 2.**
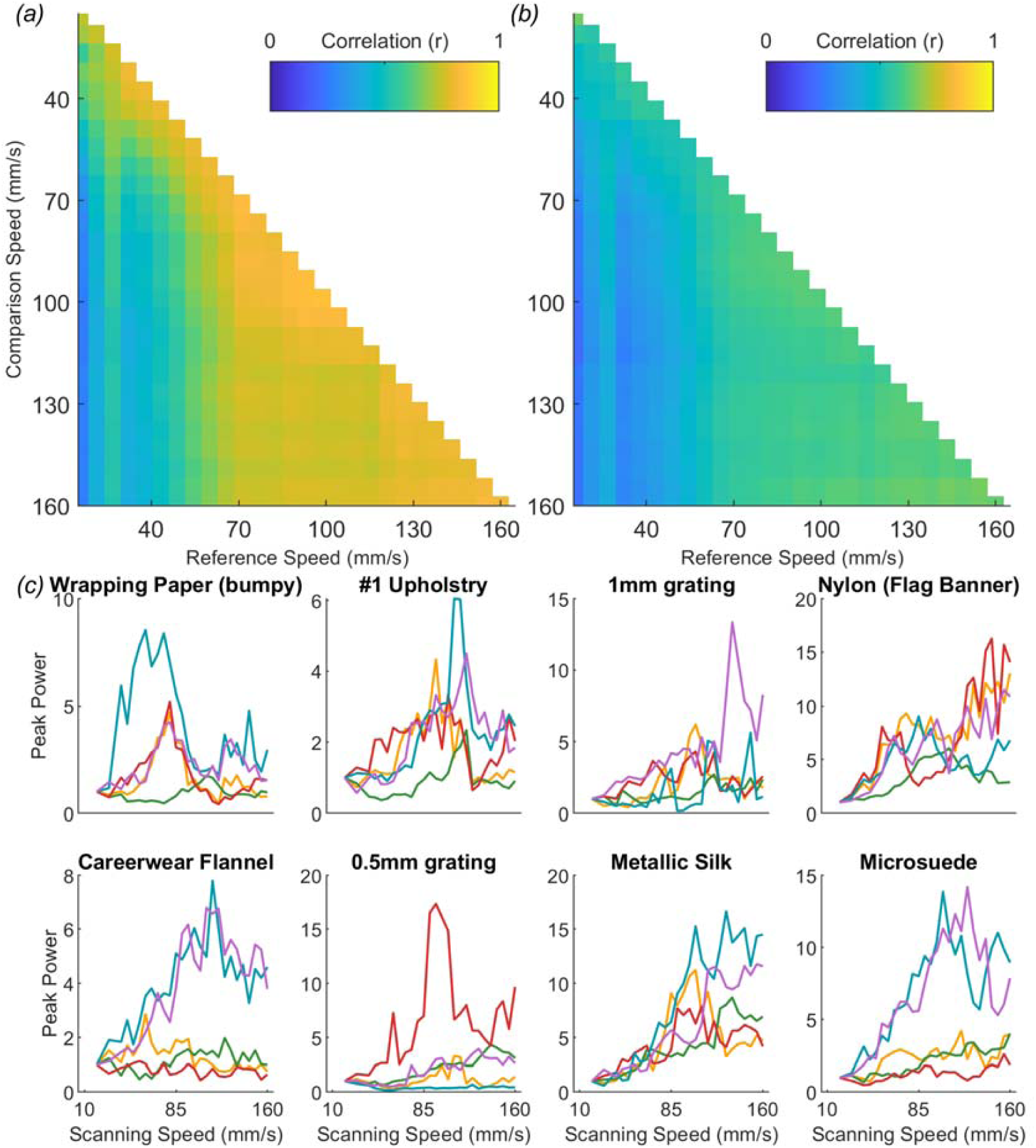
Stability of the spatial frequency composition with speed. (*a*) Average pairwise correlation within texture and participant for each speed shows that PSDs are relatively stable across speeds. (*b*) Within textures but across participants shows a similar distribution albeit with weaker correlations, suggesting that the vibrations produced by a given texture are similar across participants. (*c*) The relationship between the power at the peak frequency and scanning speed is highly dependent on texture and participant.

**Supplementary Figure 3.**
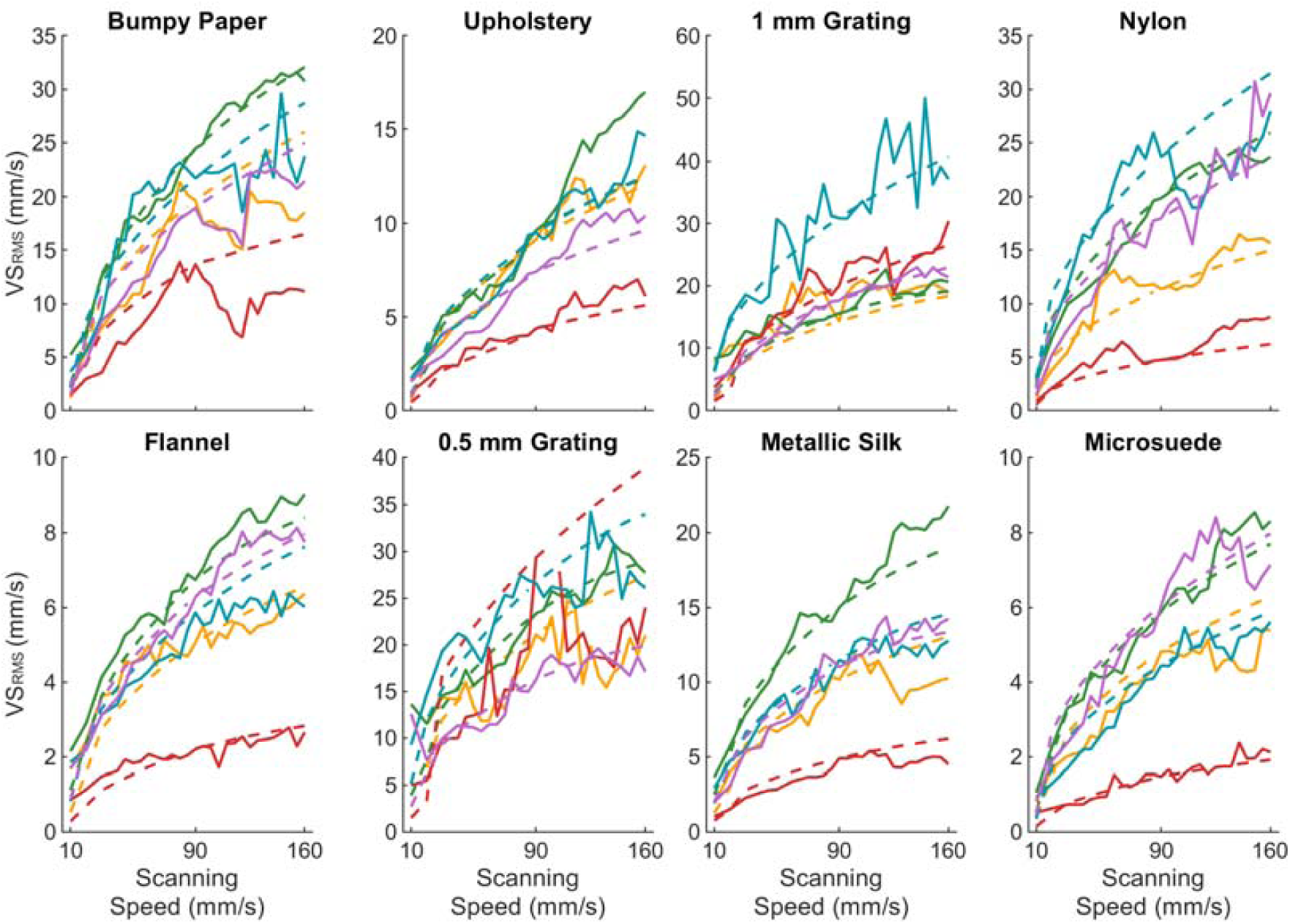
VS_RMS_ predictions based on warping the 90 mm/s trace for each texture and participant (shown in individual colors). Predictions are shown in dashed lines. The parameter-free model accurately accounts for variations across both textures and participants (each represented by a different color).

**Supplementary Figure 4.**
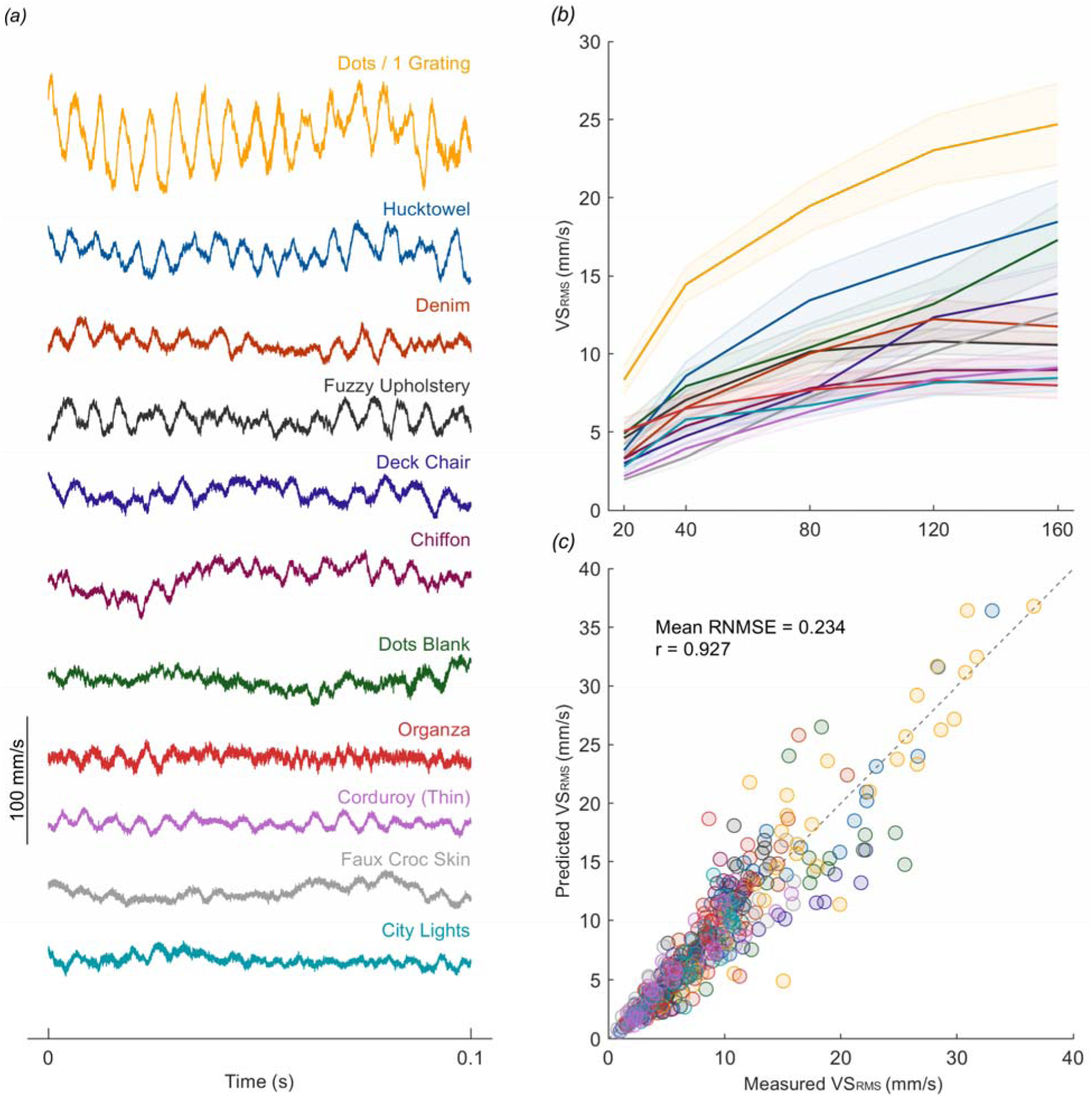
Validation of the warping model on a second dataset. (*a*) Example velocity traces of each texture at 80 mm/s scanning speed. (*b*) The RMS velocity averaged across subjects against speed for each texture. Solid line indicates mean and the shaded error the standard error. (*c*) Measured versus predicted VS_RMS_ for every texture, participant, and speed (n = 396). RNMSE is the normalized error averaged across each participant and texture, r is the correlation between all measured and predicted points.

**Supplementary Figure 5.**
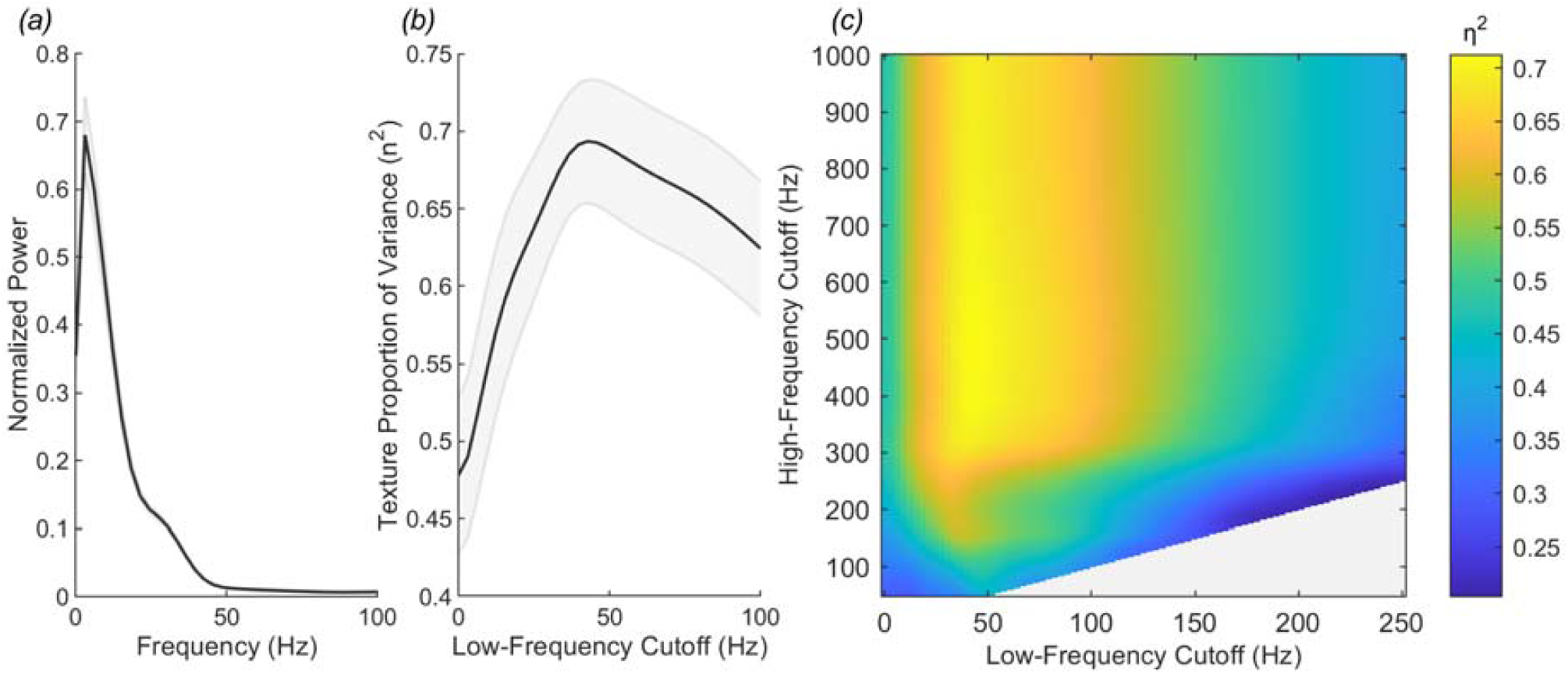
Low frequency noise dominates the signal and obscures texture related information. (*a*) Average normalized power spectral density for all traces. Much of the power is concentrated at the low frequencies (*b*) Proportion of variance attributed to texture (computed using a 2-way ANOVA) increases as the low-frequency cutoff increases up to about 50 Hz. Black line is average and gray shaded area SEM. (*c*) The average proportion of variance explained by texture across participants when changing both the low- and high-frequency cutoff.

